# Quantifying the performance of MEG source reconstruction using resting state data

**DOI:** 10.1101/248252

**Authors:** Simon Little, James Bonaiuto, Sofie S Meyer, Jose Lopez, Sven Bestmann, Gareth Barnes

## Abstract

Resting state networks measured with magnetoencephalography (MEG) form transiently stable spatio-temporal patterns in the subsecond range, and therefore fluctuate more rapidly than previously appreciated. These states populate and interact across the whole brain, are simple to record, and possess the same dynamic structure of task related changes. They therefore provide a generic, heterogeneous, and plentiful functional substrate against which to test different MEG recording and reconstruction approaches. Here we validate a non-invasive method for quantifying the resolution of different inversion assumptions under different recording regimes (with and without head-casts) based on resting state MEG. Spatio-temporally partitioning of data into self-similar periods confirmed a rich and rapidly dynamic temporal structure with a small number of regularly reoccurring states. To test the anatomical precision that could be resolved through these transient states we then inverted these data onto libraries of systematically distorted, subject specific, cortical meshes and compared the quality of the fit using Cross Validation and a Free Energy metric. This revealed which inversion scheme was able to best support the least distorted (most accurate) anatomical models. Both datasets showed an increase in model fit as anatomical models moved towards the true cortical surface. In the head-cast MEG data, the Empirical Bayesian Beamformer (EBB) algorithm showed the best mean anatomical discrimination (3.7 mm) compared with Minimum Norm / LORETA (6.0 mm) and Multiple Sparse priors (9.4 mm). This pattern was replicated in the second (conventional dataset) although with a marginally poorer prediction of the missing (cross-validated) data. Our findings suggest that the abundant resting state data now commonly available could be used to refine and validate MEG source reconstruction methods or recording paradigms.

## Introduction

Resting state network analyses have emerged as a powerful tool for deconstructing and mapping large scale networks in the human brain. These have been shown to be linked to a strikingly wide range of cognitive functions in health and neurological and psychiatric disorders (Barttfeld et al., 2015; Kaiser et al., 2015; Lewis et al., 2009; Li et al., 2012; Peterson et al., 2014; Philippi et al., 2015; Reineberg et al., 2015; Sheline and Raichle, 2013; Tessitore et al., 2012; Venkataraman et al., 2012; Wu et al., 2014; Wurina et al., 2012). Emerging evidence from magnetoencephalography (MEG) suggests that resting state networks may have a significantly finer temporal scale than previously recognised, with network state fluctuations occurring on the scale of 100-200 milliseconds (Baker et al., 2014; Koenig et al., 2002; Wackermann et al., 1993; Woolrich et al., 2013). These states are thought to be relevant because they effectively rehearse the transient dynamic patterns observed during task performance (O’Neill et al., 2017).

MEG detects electromagnetic fields at sensors outside the head with the resultant challenge of inferring the neuronal sources responsible for these measured external electromagnetic changes. One solution is to restrict the multiple potential solutions to this challenge through *a priori* assumptions relating to the model of neural activity, including the anatomical structure of the brain (the cortical mesh) and temporal relationship between sources (i.e. source co-variance matrix). Both these priors have uncertainty attached to them: for the anatomy, the uncertainty regarding head position, due to co-registration error and within-session head movement (Bonaiuto et al., 2018; Hillebrand and Barnes, 2011; Meyer et al., 2017b; Troebinger et al., 2014a, 2014b); whereas assumptions about the temporal relationship between sources are continually being refined and debated (Baillet, 2015; Baillet et al., 2001; Wipf and Nagarajan, 2009).

Here we leverage new analytic techniques to quantify the sensitivity of MEG source inversion schemes by progressively deforming the anatomical models (López et al., 2017, 2012; Stevenson et al., 2014). Specifically, here we quantify how distortions in the MRI-extracted cortical manifold affect our ability to predict or model the underlying current distribution (through cross validation error and Free Energy respectively). The rationale is that the best MEG inversion scheme will be the most sensitive to subtle distortions of the anatomy (as we know that MEG data derives from grey matter structure). This spatial distortion metric then provides a principled choice between different *a priori* inversion assumptions (i.e. different algorithms) and recording techniques.

In addition to distinguishing between algorithms, here we also tested whether we could use the same methods to distinguish between datasets collected with a head-cast (Meyer et al., 2017a; Troebinger et al., 2014a, 2014b), where the forward model is more precisely known, and those collected without.

The paper proceeds as follows. We first parcellated our two resting state datasets into brief epochs, based on the dominant spatio-temporal network during short temporal epochs, using a hidden Markov model. The epochs for the four most dominant networks were then amalgamated into four enriched datasets and taken forwards for inversion. These datasets were then inverted onto a library of subject specific distorted meshes, for which we had control over the spatial detail available in the forward model. For each of these meshes, and inversion scheme we quantified model fit using Cross validation and Free Energy metrics. We found an approximately monotonic improvement in model fit with decreasing amounts of distortion towards the real mesh. We then used this spatial quantification to compare different inversion co-variance prior assumptions as implemented in a number of different, commonly utilised schemes. For these data we found that the beamformer based priors (EBB) were the most sensitive to small deviations from the true anatomy. Finally we directly compared data recorded with head-casts to data recorded conventionally and found marginal (but not significant) differences between the two recording techniques.

## Methods

### MRI

Subjects underwent two MRI scans using a Siemens Tim Trio 3 T system (Erlangen, Germany). For the head-cast scan, the acquisition time was 3 min 42 s, in addition to 45 s for the localizer sequence. The sequence implemented was a radiofrequency (RF) and gradient spoiled T1 weighted 3D fast low angle shot (FLASH) sequence with image resolution 1 mm^3^ (1 mm slice thickness), field-of view set to 256, 256, and 192 mm along the phase (A–P), read (H–F), and partition (R–L; second 3D phase encoding direction) directions respectively. A single shot, high readout bandwidth (425 Hz/pixel) and minimum echo time (2.25ms) was used. This sequence was optimised to preserve head and scalp structure (as opposed to brain structure). Repetition time was set to 7.96 ms and excitation flip angle set to 12° to ensure sufficient SNR. A partial Fourier (factor 6/8) acquisition was used in each phase-encoded direction to accelerate acquisition. For the anatomical scan later used to construct the cortical model, multiple parameter maps (MPM) were acquired to optimise spatial resolution of the brain image (to 0.8 mm). The sequence comprised three multi-echo 3D FLASH (fast low angle shot) scans, one RF transmit field map and one static magnetic (B0) field map scan (Weiskopf et al., 2013).

### Head-cast construction

Scalp surfaces from the head-cast MRI data were extracted using SPM12 (http://www.fil.ion.ucl.ac.uk/spm/) by registering MRI images to a tissue probability map which classified voxels according to tissue makeup (e.g. skull, skin, grey matter etc.). The skin tissue probability map was transformed into a surface using the ‘isosurface’ function in MATLAB^®^ and then into standard template library format with the outlines of three fiducial coils digitally placed at conventional sites (left/right pre-auricular & nasion). Next, a positive head model was printed using a Zcorp 3D printer (600 x 540 dots per inch resolution) and this model placed inside a replica dewar-helmet with liquid resin poured between the two, resulting in a flexible, subject specific, foam head-cast with fiducial indentations in MRI-defined locations (Meyer et al., 2017a).

### MEG recording

Resting state data was acquired from 12 healthy subjects using head-casts (age: 26.6 ± 1.0 yrs) and 12 other healthy subjects without head-casts (age: 25.2 ± 1.9 yrs). All subjects were right handed, had normal or corrected-to-normal vision, and had no history of neurological or psychiatric disease. Informed written consent was given by all subjects and recordings were carried out after obtaining ethical approval from the University College London ethics committee (ref. number 3090/001).

All subjects underwent a 10 minutes resting state scan with eyes open, using a CTF 275 Omega MEG system. The head was localised using the three head-cast-embedded fiducials (head-cast subjects) or fiducials placed on the nasion and left/right pre-auricular points (non-head-cast subjects). Average absolute range of head movement within the 10 minute resting state recording was 0.26 ± 0.06, 0.24 ± 0.05, 1.1 ± 0.54 mm (X,Y,Z directions; ± SEM) for head-cast and 3.2 ± 0.5, 3.0 ± 0.5, 3.3 ± 0.2 (X,Y,Z directions; ± SEM) for non-head-cast data. The data were sampled at a rate of 1200 Hz, imported into SPM12 and filtered (4^th^ order butterworth bandpass filter: 1-90 Hz, 4^th^ order butterworth bandstop filter 48-52 Hz) and downsampled to 250 Hz.

In order to parcellate the data into self-similar periods that capture the resting state network transitions (100-200 ms), a Hidden Markov Model (HMM) was used that could identify the rapid formation and dissolution of recurring resting state networks. With this, a ‘statepath’ was estimated for each 10 minute resting state block, which tracks the fine spatiotemporal dynamics and allocates each point in time to one of eight dominant network states (Baker et al., 2014). For this statepath determination, a copy of each subject data was dimensionally reduced using principle component analysis (PCA) to derive 40 components of unit variance and mean (Woolrich et al., 2013). With these data, an 8 state Hidden Markov model (HMM; www.fmrib.ox.ac.uk/~woolrich/HMMtoolbox) was then applied to derive the most probable state at all points in time (the statepath) (Fig. 1A) (Baker et al., 2014). The continuous 600s of data were then epoched into 200ms blocks/epochs (the average scale of individual state transitions (Baker et al., 2014) and each 200ms data epoch allocated to a new dataset according to the state that was most dominant during that time period. This resulted in 8 new datasets each with a collection of epochs that corresponded to a distinct network state as identified by the HMM. The four most dominant states for each individual subject (as measured by most time spent in that state) was taken forward for further analysis and spatial estimates averaged across those four inversions for each subject. As the HMM was performed on individual, rather than group concatenated data, state numbers did not directly correspond across subjects. This however permits superior partitioning within subjects since it allows the model to optimally fit states to the individual subjects, rather than fitting individual data to group states that are common across all subjects. This resulted in 815 ± 56.9 data segments per partitioned dataset - equivalent to 163s of continuous data. Since the HMM selected periods of self-similarity within the resting state, partial correlation maps were examined for all subjects to check that segregation into separate networks had occurred and this segregation did not relate to eye blinks, muscle artefacts or cardiac interference (Figure 1B).

**Figure 1.**
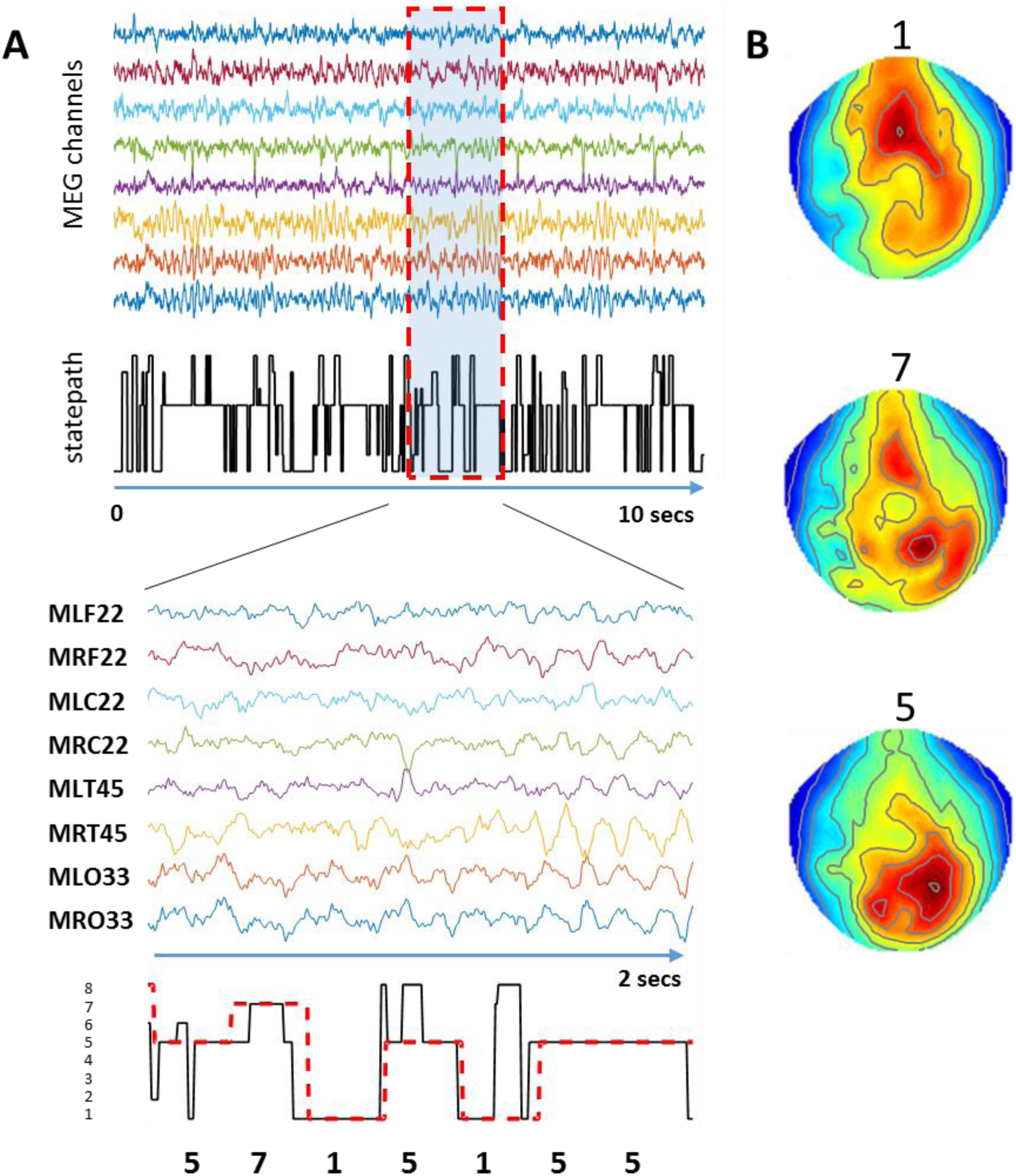
MEG data and Hidden Markov Model network allocation. **A.** Top panels show a selection of MEG sensor time series for head-cast subject 1 (representative and chosen at random). Middle panel demonstrates the statepath as determined by the HMM, showing rapid transitions between different states. Timecourses recorded from a random subset of sensors is shown. Bottom panel shows a short (2s) period of data, expanded with the corresponding section of the statepath (black line). Overlying the statepath trace (black) is the modal statepath for each 200ms period (dash red line) and this modal statepath trace was used to sort each 200ms data epoch into to 1 of 8 states according to which state was most frequent during that period. These epochs of data for each of these states were aggregated together to form 8 new datasets. **B.** The statepath traces for the three most dominant states here (1, 7, 5) are shown correlated against the time series amplitudes in all sensors to derive a map of activity (red, greater positive correlation, blue, less positive correlation) associated with each particular state. Data are shown in sensor space in order to check that differential network parcellation had indeed occurred according to the statepath detection method and was not artefactual (e.g. that the topography resembles that expected for an eye blink). In the example shown here – state 5 is the most common state – corresponding to the posterior alpha rhythm.

### Subject specific cortical mesh libraries

To extract the cortical pial mesh surface, Freesurfer software was used, optimised for MPM scans (anisotropic Freesurfer filter and a hard white matter threshold with no normalisation) (Lutti et al., 2014). Then, for each individual subject, the pial cortical mesh was taken and deformed using a 3D weighted Fourier analysis that effectively decomposes the original 3D mesh structure into spatial harmonic components (Chung et al., 2007). These are then sequentially combined to form a set of meshes of progressively increasing spatial detail (Figure 2B) (Stevenson et al., 2014). These meshes are called the Weighted Fourier Series (WFS) and result in a library of meshes for each subjects with different levels of spatial detail, from a completely smooth pair of ovoid surfaces (mesh 1) up to a mesh similar to that of the real brain (mesh 50) (see Figure 2B). The WFS can be expressed as follows:

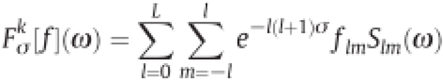

where σ is the bandwidth of the smoothing kernel (set at 0.0001), *L* is the harmonic order of the surface, *S_lm_* is the spherical harmonic of degree l and order m, and the Fourier coefficients are given by *f_lm_*=〈*f*, *S_lm_*〉, where f is determined by solving a system of linear equations (Chung et al., 2007). All meshes, including the true mesh, were downsampled by a factor of 10 in Freesurfer to ~ 33,000 vertices to aid computational efficiency.

### Source reconstruction

HMM parcellated datasets were projected from 272 spatial (3 channels damaged) and 16 temporal modes and inverted sequentially using our library of individualised and distorted cortical meshes (WFS) for each subject. Notably, using a cortical mesh further constrains solutions to those which are located at the cortical surface. We firstly implemented an Empirical Bayesian Beamformer (EBB) inversion. Beamforming makes a direct estimate of source covariance based on the assumption that there are no zero-lag correlated sources, a form of spatial filtering that ensures that no two parts of the brain has exactly the same neuroelectric activity at any given time. This was compared with alternative inversion schemes that are based on different prior assumptions regarding source co-variance. Here we used standard (and non-optimized) versions of Minimum norm (MMN) (Uutela et al., 1999), LORETA (LOR) (Pascual-Marqui et al., 2002), Empirical Bayes Beamformer (EBB) and Multiple Sparse priors (MSP) (Friston et al., 2008) inversion schemes as implemented in SPM12. Each of these methods can be described by a different selection of source covariance matrices (Belardinelli et al., 2012; Friston et al., 2008; López et al., 2014).

In order to quantify the quality of the fit we used two complementary metrics: cross validation and Free energy. For cross validation error, a subset of sensors (10%; 27 sensors) are turned off. An estimate of current flow is made based on the remaining 90% of the MEG channels (and a specific inversion scheme). This current flow is then projected back outside of the head to the 10% of disabled sensors and the error between the predicted and measured data calculated. This was repeated 10 times for each inversion with different randomly selected subsets of removed sensors for each iteration. The cross validation error was then converted to a percentage of source data explained and averaged across all iterations and across the 4 network datasets inverted per subject. The Free Energy metric approximates the log model evidence for the final generative model based on all of the sensor data by rewarding the accuracy of fit whilst also penalizing model complexity (Friston et al., 2007).

Both cross validation error explained and Free Energy provide metrics which can be used to directly compare different models of the same data with increasing values of each suggesting an improved model fit (Henson et al., 2009). Whilst the Free Energy provides a useful relative model fit metric for any given dataset the absolute value is data dependent and therefore cannot be used to compare between datasets. In contrast, the cross validation error explained gives a meaningful quantification of the total amount of data explained and can be used to compare across the different groups (head-cast versus non head-cast).

Here we performed these inversions using our subject specific libraries of anatomically degraded meshes from the WFS with different levels of distortion and compared them to an inversion performed using the real mesh for each subject. In order to facilitate comparisons of how the anatomical distortion affected the inversions for different subjects with different baseline measures of model fit to their real brain meshes, we normalised the cross validation percentage error explained and free energy metrics by subtracting the values for the real mesh to derive a relative measure (ΔCV & ΔF). As such, a worse fit gives a negative value of ΔCV / ΔF and we would predict that as the anatomical complexity of the mesh increases (mesh is less distorted) and approaches that of the real mesh – the quality of the model should improve and the ΔCV / ΔF should approach zero. Across our group of subjects we determined the level of distortion at which this first becomes statistically distinguishable from the real mesh using a t-test of ΔCV values for each harmonic compared to zero and similarly for ΔF using Bayesian model comparison. This point, labelled the highest distinguishable harmonic (HDH), identifies the minimum amount of mesh distortion that can be reliably distinguished from the real mesh by inversion and can be converted into a conventional spatial metric (mms) by comparing vertex distances. The three dimensional (Euclidean) distance (in mm) between every vertex from the HDH to the corresponding vertex on the true mesh was therefore calculated and the upper, 95th percentile, distance averaged over all vertices in the mesh. This method is more conservative than that previously employed (Stevenson et al., 2014), as this directly matches corresponding vertices and therefore reduces the underestimation that could result if, following harmonic distortion, a vertex now lies closer to a non-corresponding other vertex. This distance therefore represents an upper bound on the spatial discriminability of both head-cast and non-head-cast resting state data.

### Control analyses

In order to verify our findings, we performed a number of control analyses with distorted data/sensors positions for the head-cast / EBB inversion. Firstly, we used our same data but destroyed its correspondence with the MEG sensor locations by randomly shuffling the MEG channels labels and repeated the analysis above 10 times for each subject and averaged over the ΔCV and ΔF for these different shuffled dataset inversions. Following this, we used the correct sensor labels but next degraded our data by introducing different amounts of scaled white noise to change the signal-to-noise ratio of the sensor level data (5 dB TO - 20 dB) (Troebinger et al., 2014a). In both cases (sensor shuffling and noise addition) one would expect the ability to discriminate the true generative model from distorted ones to decrease.

## RESULTS

### Anatomical cortical model

In order to determine the sensitivity of the inversion to the level of detail in the underlying anatomical mesh model we calculated the ΔCV and ΔF for each subject in the head-cast dataset across their subject specific library of distorted meshes (Fig. 2). This showed increasing cross validation sensor data explained (ΔCV; reduced error) and increasing Free energy (ΔF) for all subjects. Statistical group level testing revealed that meshes lower (more deformed) than the 35 harmonic could be distinguished from the real mesh by ΔCV (HDH 35; t_11_= −2.49, p=0.03) and lower than 31 by ΔF (BMC, exceedance p = 0.046; Fig. 2)

**Figure 2.**
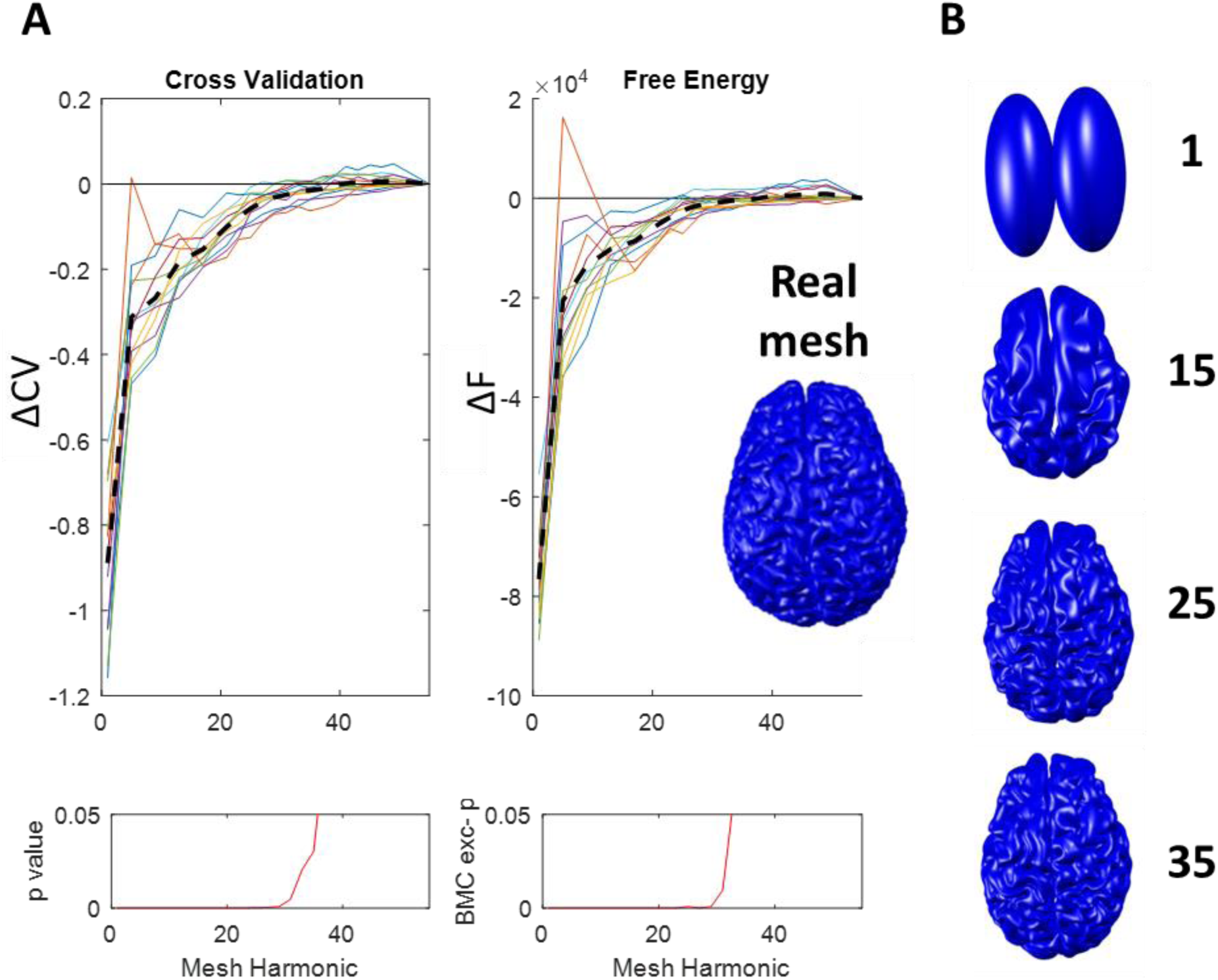
Relative Cross Validation and Free Energy results for a library of different meshes for head-cast resting data using an EBB inversion. **A.** Increasing cross validation data explained (ΔCV) and relative Free Energy ΔF with improving spatial resolution of harmonic meshes (from left **→** right). Top plots show individual subject ΔCV (left) and ΔF (right) values with superimposed mean value (dashed black line). Lower panels show the group level statistical significance using t test (ΔCV) and Bayesian model comparison (ΔF). **B.** Example selection of meshes from 1 subject showing different levels of distortion, from smooth ovoid surfaces (WFS mesh 1) up to real mesh (shown inset).

### Inversion algorithms / source covariance priors

The effect of source co-variance prior assumptions was then assessed by repeating the process for three other commonly implemented inversion algorithms (MNM, LOR, MSP) on our head-cast dataset. These showed lower anatomical mesh discriminability for all alternative algorithms with an HDH for MNM of 25 (t_11_=−1.80 -, p=0.036), an HDH of 25 (t_11_=−1.79; p=0.039) for LOR and an HDH of 17 for MSP (t_11_=−2.55; p=0.027) (Fig 3A).

The mean distance between vertices on these meshes and the real mesh was then calculated for each subject. These were then averaged to give a spatial measure of anatomical discriminability (Fig 3A), under the different prior covariance assumptions as implemented in the different inversion algorithms. This ranged from 3.7 mm for EBB to 6.0 mm for MNN/LOR and 9.4 mm for MSP (Fig 3B). Directly comparing the absolute cross validation error explained (CVp) across inversion conditions (using the real mesh) demonstrated a significant difference between EBB and the other 3 algorithms (EBB/MMN; paired t-test – t_11_=7.6, p<0.001; EBB/LOR; paired t-test – t_11_=7.6, p<0.001; EBB/MSP; paired t-test – t_11_=14.4, p<0.001).

**Figure 3.**
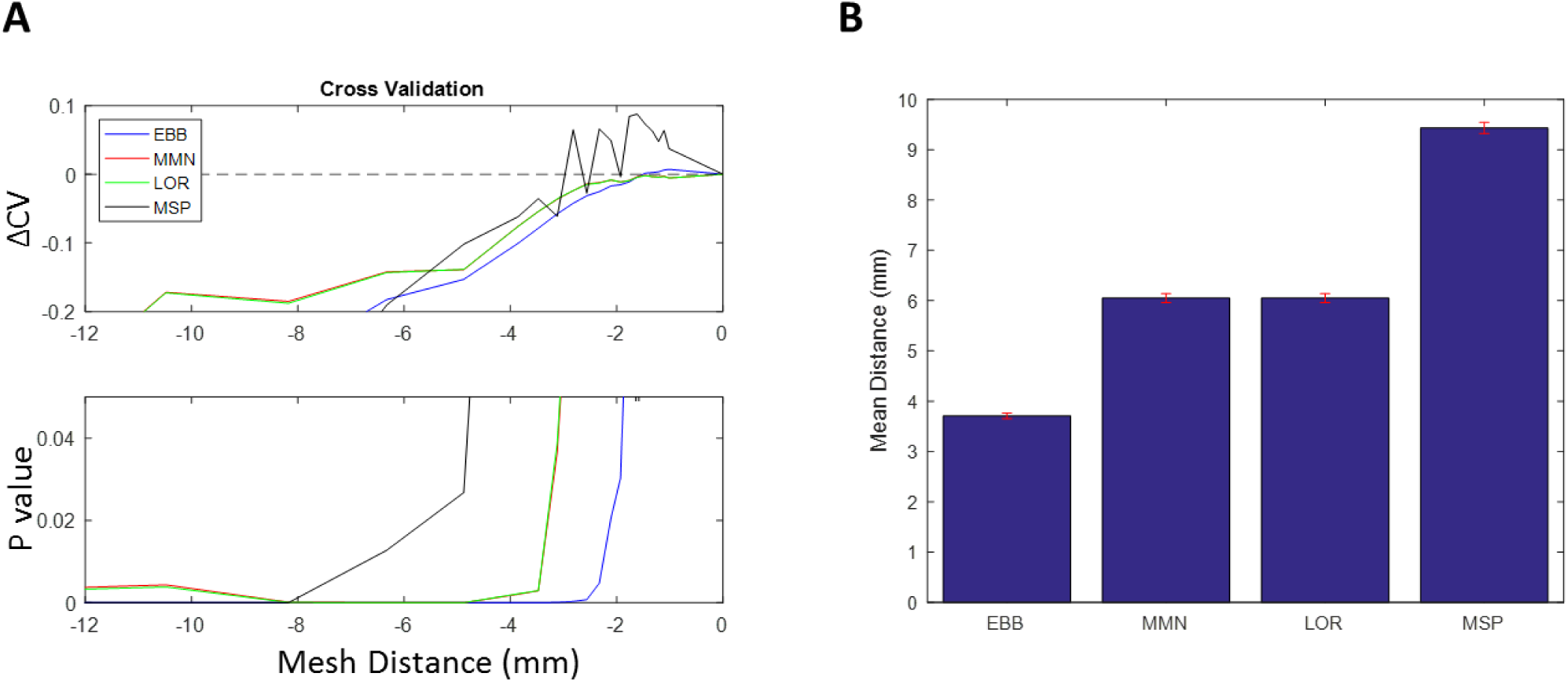
Effect of different inversion schemes on head-cast resting state resolution. **A.** Relative cross validation error explained (ΔCV) for head-cast MEG dataset (12 subjects), shown for 4 different inversion types (EBB, MNM, LOR & MSP) according to mean distance of the distorted mesh (x axis) from the real mesh. Lower panel shows the group level statistical significance by t-testing (note MMN and LOR have very similar values and therefore lines are closely overlapping). **B.** Mean distance of vertices from highest significant mesh identified in the left hand panel and the real mesh for each subject. Error bars show SEM of this distance across subjects using their individualised meshes and group level HDH.

Notably – a 2 factor within subject ANOVA of cross validation with factors – inversion type (EBB, MMN, MSP) and mesh smootheness (harmonic 1 & harmonic 50) showed a strongly significant interaction between inversion type and mesh harmonic level (F_2_=29.9, p<0.000). Post hoc examination showed that this was driven by a stronger effect of mesh distortion on the MSP inversion algorithm than EBB or MMN.

### Head-cast versus conventional MEG

The EBB algorithm was therefore taken forwards for a comparison of head-cast versus non – head-cast datasets (Fig 4). We found that the HDH was higher for the head-cast recorded dataset at 35 (t_11_= −2.49, p=0.03) than for the non – head-cast related dataset at 29 (t_11_=−2.70, p=0.021). Furthermore, the absolute (non-normalised) cross validation error explained (CV) was higher in the head-cast (83.2 ± 0.71 %) than in the conventional recordings (79.7 ± 1.79 %) although this did not reach significance (t_11_=1.79; p=0.1).

**Figure 4.**
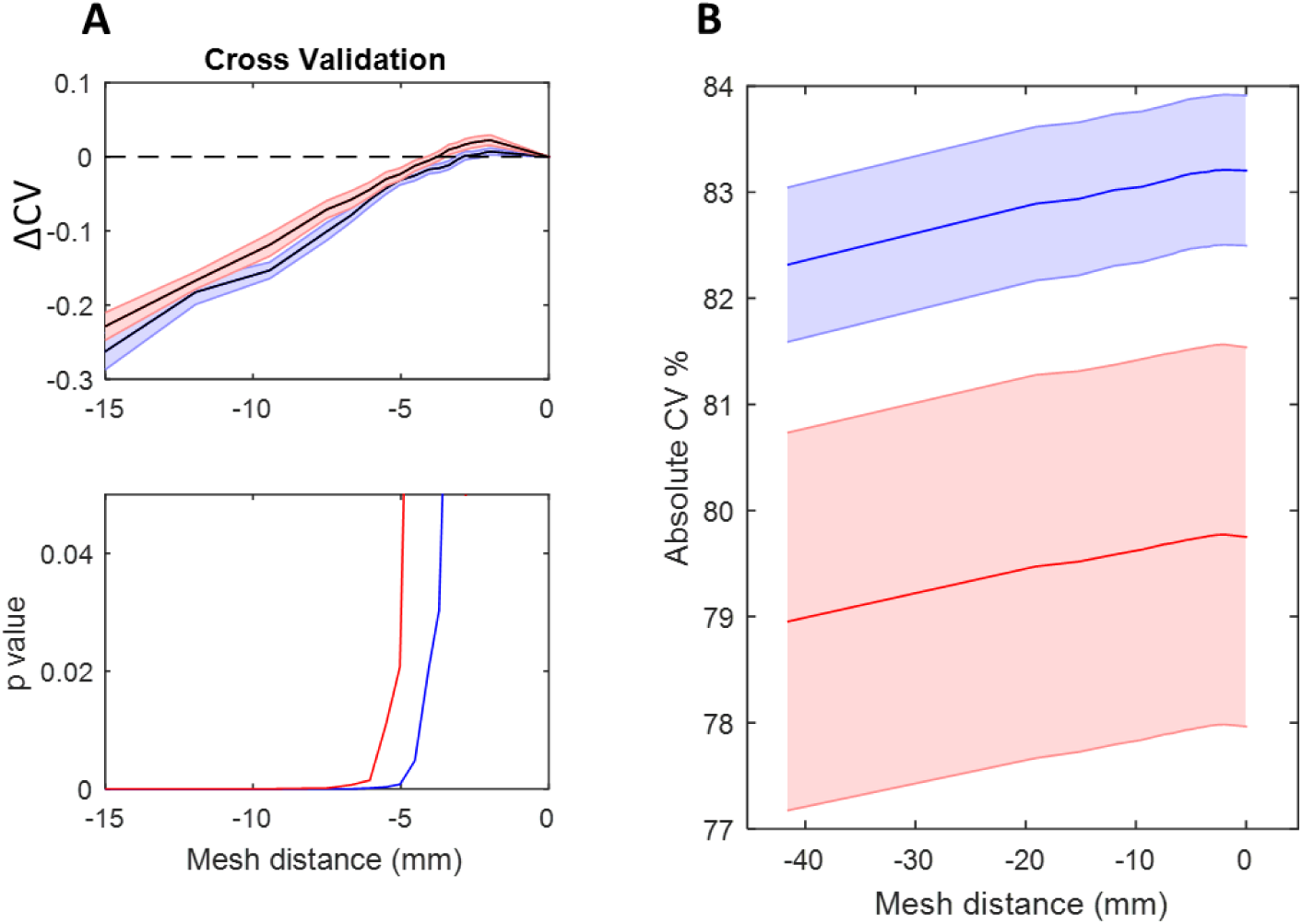
Comparison of head-cast versus conventional MEG Cross validation results. **A.** Comparison of ΔCV for different recording methodologies with head-cast data (blue) showing higher discrimination (35) than non-head-cast data (red; 29). **B.** Absolute cross validation error explained is also higher in the head-cast versus the conventional MEG. Note that this holds for all inversion types and for all levels of mesh distortion, but was not significantly different on statistical testing.

### Control analyses

Finally, we checked our analyses by repeating our inversions (Head-cast dataset, EBB algorithm) but only after degrading the consistent relationship between our sensor positions and sources by shuffling the sensor labels. As expected, this resulted in a breakdown of the previously shown relationship between Cross validation and Free Energy with mesh distortion. Notably we found that lower harmonics (smoother) meshes now showed higher ΔCV (Fig 5A) (smoother surfaces superior when sensors shuffled). Thereafter we degraded our data (without sensor shuffling) by the addition of varying levels of Gaussian white noise (5 → −50 dB) and again repeated our analysis (Figure 4B). This demonstrated that with increasing levels of noise (decreasing SNR), the curve describing the relationship between Cross validation and harmonic mesh function flattened and the crossing point (HDH) reduced, indicating that the inversion was no longer able to statistically distinguish the more complex meshes from the real mesh, as would be expected if the data are primarily noise.

**Figure 5.**
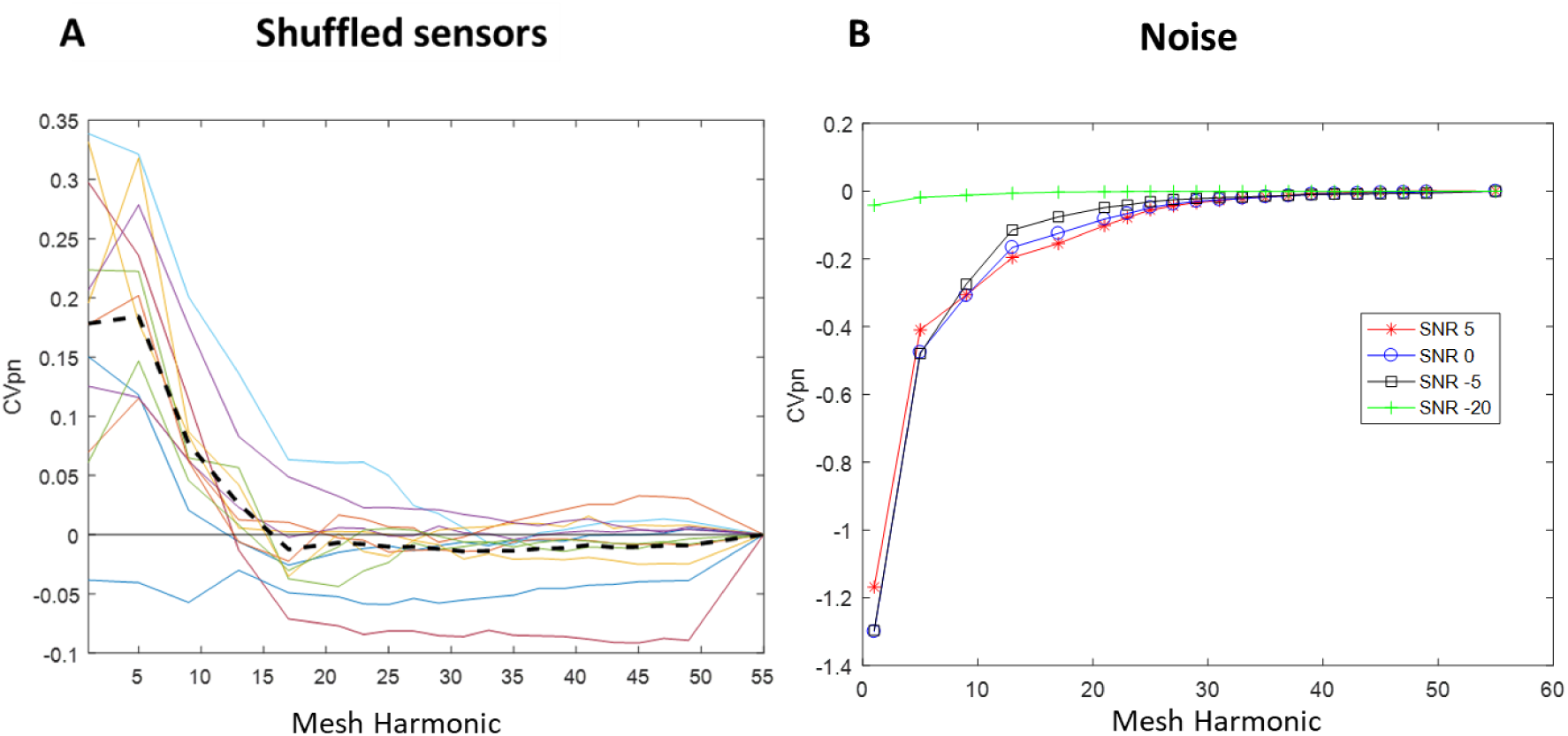
Effects of shuffling sensors (A) and replacing data with varying levels of Gaussian White noise (B). **A.** The relationship between the Cross validation % error explained (ΔCV) of inversion and increasing harmonic mesh after random shuffling of sensor labels. Note the decreasing Cross validation fit with increasing mesh harmonic (increasing mesh detail) with the dashed line showing the average across all subjects. **B.** The relationship between (mean of all 12 subjects) ΔCV of inversion and increasing harmonic mesh after addition of Gaussian white noise across a range of noise additions (from SNR 5 to SNR −20).

## Discussion

Resting state data is rapidly dynamic, emulates task induced network changes and is simple to acquire (Baker et al., 2014; O’Neill et al., 2017). Here we show that it can provide a ready substrate for principled testing of MEG recording methods and inversion assumptions including anatomical forward modelling and functional (co-variance) priors.

We showed that in moving the cortical surface from heavily distorted to the true anatomy, all of the models, based on commonly used imaging assumptions, showed a significant and monotonic improvement. This improvement saturated for some imaging assumptions before others, with the beamformer based algorithms (closely followed by Minimum norm) continuing to improve up until the cortical surface deviated by on average 4mm from the ground-truth. We also found that a marginal, although non-significant, improvement in our models when using MEG data based on head-cast recordings compared to conventional recordings. Critically rather than compare methods through simulation or through a limited task set (with ground truth from another modality) we have presented a method to optimize MEG recording methods, forward and inverse models without introducing selection bias and based on a plentiful supply of non-invasive human data.

We compared between algorithms in two ways: by the model fit (or amount of data predicted), and by comparing the sensitivity of each algorithm to the true anatomy. These two tests need not necessarily have been in accord. For example- had we used a bunny-shaped blancmange mould instead of a cortical surface we would still have been able to rank the algorithms based the amount of data predicted; but we would not have expected any monotonic improvement as features were added to the bunny. A related control analysis (figure 5) is that when used the same data but with shuffled lead-fields (destroying the link between the sensors and the anatomy) the amount of data we are able to predict actually decreases as the cortical model approaches the truth. It is therefore striking that the models that benefitted most from the true cortical manifold were also those that predicted the most data. This not only adds anatomical validity (confirming that the data being described is indeed generated by pyramidal cell populations normal to the cortical surface) but also allows us to quantify algorithm performance in millimetres (Stevenson et al., 2014).

Across anatomy (Fig. 2) and inversion assumptions (Fig. 3), the parametric (Free energy from empirical Bayes) and non-parametric (cross-validation) metrics of model fit were in accord. This helps build confidence in the parametric free-energy metric which is considerably faster, makes use of all the available data, and has a direct probabilistic interpretation. The Bayesian formalism is however predicated on comparing how different models explain the same data; the use of cross-validation, which provides an absolute quantitative measure of data predicted, also allowed us to compare between different datasets (head-cast and non-head-cast).

Resting state MEG analysis has been impeded by the spectral/dimensional complexity of the datasets (Vidaurre et al., 2016) as well as reduced spatial resolution that limits the ability to discriminate different sources (Colclough et al., 2016; Liuzzi et al., 2016). Recent improvements in methods have permitted enhanced temporal discrimination (Baker et al., 2014; Woolrich et al., 2013). These and other development have resulted in the emergence of a number of potential clinical applications for resting state MEG, although these have yet to transition to clinical utilisation, in part due to reduced spatial resolution (Bosboom et al., 2006; Bosma et al., 2009; Hinkley et al., 2011). It is notable that most of the empirical MEG literature on the resting state is dominated by what we found here to be the most likely functional priors (beamformer and Minimum norm assumptions).

We confirm here earlier reports that within-session head movements are greatly reduced for head-cast versus non-head-cast MEG (Bonaiuto et al., 2017b; Meyer et al., 2017a, 2017b) however we were surprised that the head-cast did not offer a greater modelling improvement over the non-head-cast data. Empirically this could be due to recording problems- for example in some subjects it is possible that their heads were not within the head-casts in their expected position. i.e. although the head-casts remained still the subject’s anatomy was not where we expected it to be. Another limitation could be that the models we are using do not capture the physics or physiology of the generators of the measured magnetic fields; and that the resolution is constrained by the models and not the recording. This could include for example, unmodelled noise sources such as the heartbeat, eye-blinks and other sources of noise. Although the HMM states we used were visually inspected to avoid common artefacts such as eye-blinks, it is possible that some of the modelling deficiencies come from failure to explain data that does not arise from the cortex (eg heart-beats, passing cars etc). Finally, the head-cast and non-head-cast cohorts did not overlap and a more sensitive analysis would have been to perform a within subject comparison.

We found the Multiple Sparse Priors algorithm had the least dependence on the true anatomy and also explained the least data. We should note however that the MSP-based analyses implemented here were generic and constructed from a limited set of 512 patches (or priors) placed at evenly spaced vertices. The MSP algorithm, although perhaps the most elegant and comprehensive method we tested, is also computationally disadvantaged by the need to search over a large space of possible patch/prior combinations and the inherent pitfalls of local extrema in this optimization. A more robust way to implement this algorithm would have been to select the best model from many random patch choices (Troebinger et al., 2014b).

This study was analysed using broadband (1-90 Hz) resting state data. Therefore, whether a similar level of spatial discriminability at the mm scale can be demonstrated when more selective data is used (e.g. frequency filtered or spatially restricted) remains to be shown. For example, future work might test different frequency bands (eg <30Hz, >30Hz) against different anatomy for example infra/supra granular cortical surfaces (Arnal and Giraud, 2012; Bastos et al., 2012; Bonaiuto et al., 2017a, 2017b).

## Conclusion

This study uses resting state data to compare different forward models, inversion assumptions and recording methods to provide a principled (non – biased) method for optimising and quantifying source localisation. All source localisation techniques were able to benefit from increasing anatomical precision in the underlying model but this was most pronounced for EBB in head-cast recorded data. Using this method, we demonstrate a notably high sensitivity (<4mm) to underlying anatomical distortion and also provide evidence for the anatomical validity of cortical sources in MEG source inversion.

